# Development and validation of novel proliferation-related gene signature for MSS colorectal cancer prognosis

**DOI:** 10.1101/2024.11.13.623354

**Authors:** Sen Wang, Zhiyu Yu, Peng Xu, Cheng Zhang

**Author notes:** Cheng Zhang and Peng Xu are corresponding authors E-mail addresses (C. Zhang), (P. Xu). Authors equally contributed to this work.

## Abstract

Colorectal cancer (CRC) is classified into microsatellite instability (MSI) and microsatellite stabilized (MSS) based on the differences in DNA repair mechanisms. In recent years, many studies have proved that some emerging therapeutic approaches have good therapeutic effects on MSI as a subtype but are ineffective for MSS as a subtype. This paper aims to start with the molecular mechanism of MSS and try to find the characteristic target genes of MSS as a subtype. We used bioinformatics techniques and machine learning to find GTF2IRD1. The gene GTF2IRD1 is highly expressed in MSS-type CRC but is stable in MSI. Subsequently, we verified its expression and function with relevant experiments. We demonstrated that GTF2IRD1 can promote the proliferation of MSS-type colorectal cancer and correlate with the poor prognosis of MSS-type colorectal cancer patients, which may provide a potential target gene for the clinical treatment of MSS-type colorectal cancer.

## 1. Introduction

Colorectal cancer (CRC) is highly prevalent(1), both in men and women(2). It is a very lethal heterogeneous cancer. One particular genetic subtype of CRC is microsatellite instability (MSI)-related tumors(3), which have a high tumor substantial immunological infiltrates and mutational burden (TMB)(4). For instance, in cancers, monoclonal antibodies used in Immunotherapy to target PD-1/PD-L1 are successful(5). For other varieties of CRC, Immunotherapy is not as successful with microsatellite stabilized (MSS) as monotherapy(6). A promising new treatment called Immunotherapy is being attempted in conjunction with additional therapeutic approaches to treat colorectal cancer of the MSS type(6, 7). However, there is still an ineffectiveness of treatment for MSS-type CRC patients(8).

In this work, we screened for differential genes that characterize MSS colorectal malignancies and evaluated the predictive value of these genes in patients with MSS-type CRC using a combination of bioinformatics tools and two machine learning algorithms(9). Upon closer examination of these genes, GTF2IRD1, which is highly expressed in MSS-type CRC but is stable in MSI(10), was discovered(11). Subsequent tests were conducted to demonstrate the function of this gene, and the results indicated that GTF2IRD1 promotes the proliferation of MSS-type colorectal cancer. In conclusion, this study provides a scientific platform for studying the probable pathophysiology of MSS colorectal cancer and gives a new checkpoint for its treatment(12).

## 2. Materials and methods

### Patients and data

Fifty samples were obtained from CRC patients who underwent surgical remedy at the General Hospital of the Northern Theater of Operations of the People’s Liberation Army (PLA) from July 2022 to March 2024. These included 30 patients with postoperative pathology showing MSS-type CRC and 20 patients with postoperative pathology showing MSI-type CRC. The data and samples obtained were anonymized. The samples were validated by histopathologists and clinicopathological data were maintained for each validated sample. Prior approval was obtained from the Medical Ethics Committee Branch of the General Hospital of the Northern Theater of Operations of the People’s Liberation Army(Y (2022)063), and all the participants signed an informed consent form. All methods were carried out in accordance with relevant guidelines and regulations. The GEO database repository was accessed on November 3, 2023, and gene expression matrices and clinical data were retrieved. GSE 44076 comprises colorectal and paracancerous tissue samples from 98 sufferers. GSE 25071, on the other hand, incorporates four regular tissue samples, 38 MSS-type CRC tissue samples, and 8 MSI-type CRC tissue samples. Download clinical and gene expression data for CRC patients from the TCGA database.

### PCA

We can examine the distribution patterns between tumor and normal samples, do more data analysis, and create a more dependable database for study using the Principal Component Analysis (PCA) method.

### Differential gene analysis

For normalization and log2 transformation, we employed the R′Bioconductor software package’s Linear Modeling of Microarray Data (LIMMA) technique. P values of 0.05 and fold changes of 1.5 were used to classify genes as differentially expressed. Differentially expressed genes (DEG) were found during data prediction using the GSE 44076 dataset. The “heat map” package (v1.0.12) in R can create heat and volcano maps for data visualization.

### Enrichment analysis

We analysed DEG for GO, KEGG and GSEA pathway enrichment using R software.

### LASSO, SVM-RFE

LASSO regression is a technique for performing dimensionality discounts to keep away from overfitting in large-scale gene expression records analysis. In Support Vector Machine - Recursive Feature Elimination (SVM-RFE), all features are used for classification. Then, the least weighted feature is removed from the feature set based on its importance(13).

### Survival analysis

DEGs had been recognized in colorectal cancer samples, and the expression information had been mixed with scientific prognostic information from people living with colorectal cancer in the TCGA dataset for survival analysis.

### Cell culture and transfection

HT29(EallBio, CA) and LOVO cell(EallBio, CA) lines were cultured with M5A culture medium containing 10% fetal bovine serum(Cellterlife., CA) and 1% penicillin/streptomycin(Cellterlife., CA). All cells were placed in a cell culture incubator with an incubation environment of 37° C and 5% CO2. The increased medium used to be modified every two days. When the cell subculture reached 80% fusion, the passaged cells had been digested with 0.25% trypsin/EDTA. Small interfering RNA (siRNA) focused on GTF2IRD1 and negative control siRNA (NC) were added into the cells for the usage of liposomes, respectively. Then, after 48h, cells were gathered for similar analysis. QRT-PCR detected the impact of si-GTF2IRD1 on every cell line.

### CCK-8

To detect the proliferation of co-cultured cells, a CCK-8 kit (Solarbio, CA) was used. Cells were first inoculated in 96-well culture plates, and 100 µL of complete culture solution was added to each well. Next, the healthy plates were placed in a cell culture incubator for 0, 24, 48, 72, and 96 hours. At the cease of incubation, the supernatant was removed, and ninety µL of sparkling basal culture solution was delivered to every well, accompanied by 10 µL of CCK-8 solution. Subsequently, the wells were incubated in a cell culture incubator for 2-6 h. Absorbance was measured at 450 nm using a Multiskan MK3 microplate instrument from ThermoFisher Science, USA. Each set of experiments consisted of 5 replicate wells.

### QRT-PCR

Firstly, the tissues were well ground and total RNA was extracted by adding TrIzol reagent (Thermofisher, USA). Thoroughly mixed and centrifuged at 12,000 g for 15 min. The removed 500 µL of supernatant was mixed thoroughly with 500 µL of isopropanol, which was dissolved in 20 µL of enzyme-free sterilized water, and the amount was quantified by microtitration using a NanoDrop Model 2000. According to the instructions, a total amount of RNA was mixed with TaqMan Reverse Transcription Reagent (Takara Bio, Japan) for a reverse transcription reaction. Quantitative PCR was performed using TB Green® Premix Ex Taq™ II (Tli RNaseH Plus). The primers used were 5 ‘-CAGCCTCGTGTCTGCCTTAG-3’ (forward) and 5 ‘-TTCCGGGCATTCAGGAACATT-3’ (reverse) for GTF2IRD1, and 5 ‘-GGAGCGAGATCCCTCCAAAAT-3’ (forward) and 5 ‘-GGCTGTTGTCATACTTCTCATGG-3’ (reverse).

### Immunohistochemistry

Anti-GTF2IRD1 antibody (Proteintech Group, China). First, paraffin was removed from the sections with xylene and gradient alcohol, dehydrated, and incubated in a 3% hydrogen peroxide solution for 15 min. Next, heat-mediated antigen repair was performed using citrate buffer. Subsequently, primary antibodies were added dropwise to the tissue sections and incubated at four °C overnight. The next day, the secondary antibody was added dropwise to the sections and reacted for 30 minutes at room temperature. After washing, DAB colour developing reagent was added dropwise and the reaction was observed under the microscope while the reaction was going on, and the reaction was terminated in time, and then hematoxylin was used for contrast staining. Finally, the sections were sealed with impartial gum, and observation of staining results by microscopy.

### Statistical analysis

We carried out statistical analyses the usage of R software. We considered genes with p < 0.05 and logFC_t> 1.5 as differential genes. And the expression of logFC_t> 1.5 was referred to as up-regulated expression, and the expression of logFC_t< 1.5 was referred to as down-regulated expression. The t-test was used to analyse typically distributed variables and the U-test to check non-normally distributed variables. When the p-value was a great deal much less than 0.05, we considered this as a signal of good-sized difference.

## 3. Results

### 3.1: Screening for MSS-type colorectal cancer-related differential genes

We divided the samples in the GSE44076 dataset into the tumor and normal groups to obtain MSS-type colorectal cancer-associated differential genes. We proposed preprocessing the two groups of data with principal component analysis. The results showed that the distribution patterns between tumor and normal samples had significant principal component differences (Fig 1a). Then, we found the platform number of GSE44076 and downloaded its probe annotation file in the platform. Then, we performed differential gene analysis and used the “dplyr” package to integrate the gene annotations into the differential gene table. The consequences confirmed that 310 genes had been drastically up-regulated and 482 genes had been extensively down-regulated in the tumor team, contrasting with the ordinary team (Fig 1b and c). We carried out an enrichment evaluation of these differential genes. GO enrichment evaluation confirmed that BP was once extra enriched in the pathways of extracellular structure organization, external encapsulating structure organization, and extracellular matrix organization.CC was once greatly enriched in the collagen-containing extracellular matrix and apical part of the cell pathway.MF used to be extra enriched in receptor-ligand activity and signaling receptor activator activity pathway(Fig 1d).KEGG enrichment evaluation confirmed that up-regulated differential genes have been generally enriched in the IL-17 signaling pathway, cell cycle, and associated pathways. In contrast, the sulfur had exceptionally enriched down-regulated differential genes (Fig 1e). GSEA enrichment evaluation confirmed that MSS up-regulated genes had been mainly enriched in the CELL cycle and other associated pathways (Fig 1f). So far, we screened out the MSS-type colorectal cancer-related differential genes and analyzed their enrichment.

**Fig 1.**
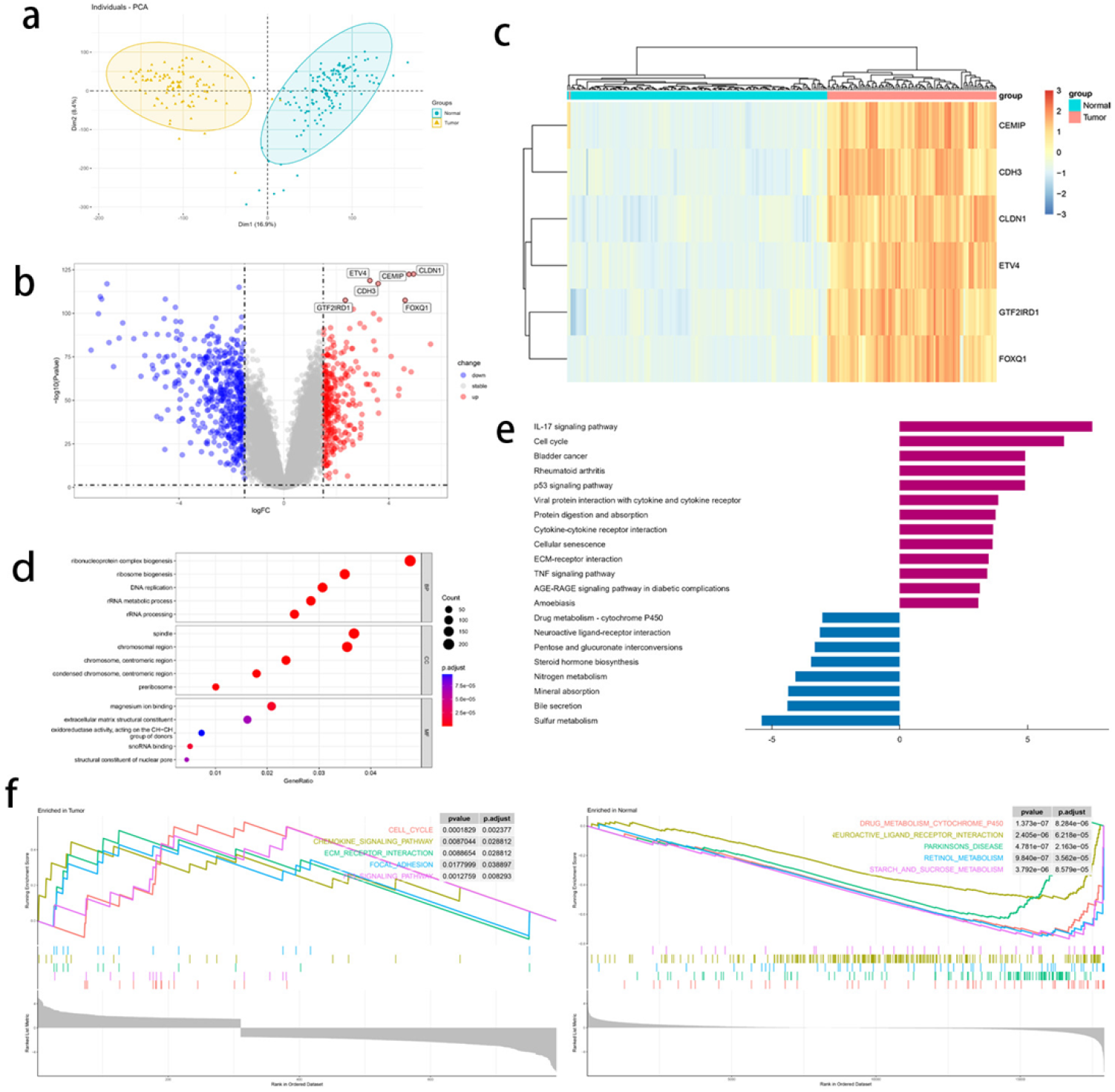
Screening for MSS-type colorectal cancer-related differential genes. (a)Principal component analysis of the two groups in GES44076. The DEG between CRC specimens and normal specimens in the GSE 44076 dataset are shown in the volcano plot of (b) and the heatmap of (c). (d) G.O. enrichment analysis. (e)KEGG enrichment analysis. (F) GESA analysis.

### 3.2: Screening for feature genes through machine learning

We used two machine learning algorithms for dimensionality reduction clustering to find valuable biomarkers for MSS-type colorectal cancer. We downscaled the screened DEGs using lasso regression, and finally, 14 genes were identified as potential diagnostic biomarkers for MSS-type colorectal cancer(Fig 2a). The SVM-RFE method was utilized, and the model was optimal when 34 features were retained, suggesting that these features may be the most important to the model’s predictive effectiveness. Therefore, in practical applications, one can consider training the model using only these 34 features to achieve better prediction performance while avoiding overfitting(Fig 2b). We obtained six overlapping genes (CLDN1, CEMIP, ETV4, CDH3, GTF2IRD1, FOXQ1)(Fig 2c). Furthermore, the results of ROC analysis confirmed the diagnostic value of these six key genes in predicting MSS-type colorectal cancer (Fig. 2d). Subsequently, we did ssGSEA analysis to affirm the interaction between these characterized genes and immune cell infiltration, and the results showed that these genes reduced most of the immune cell infiltration(Fig 2e). We found the MSS-type colorectal cancer signature genes by down-clustering the above differential genes.

**Fig 2.**
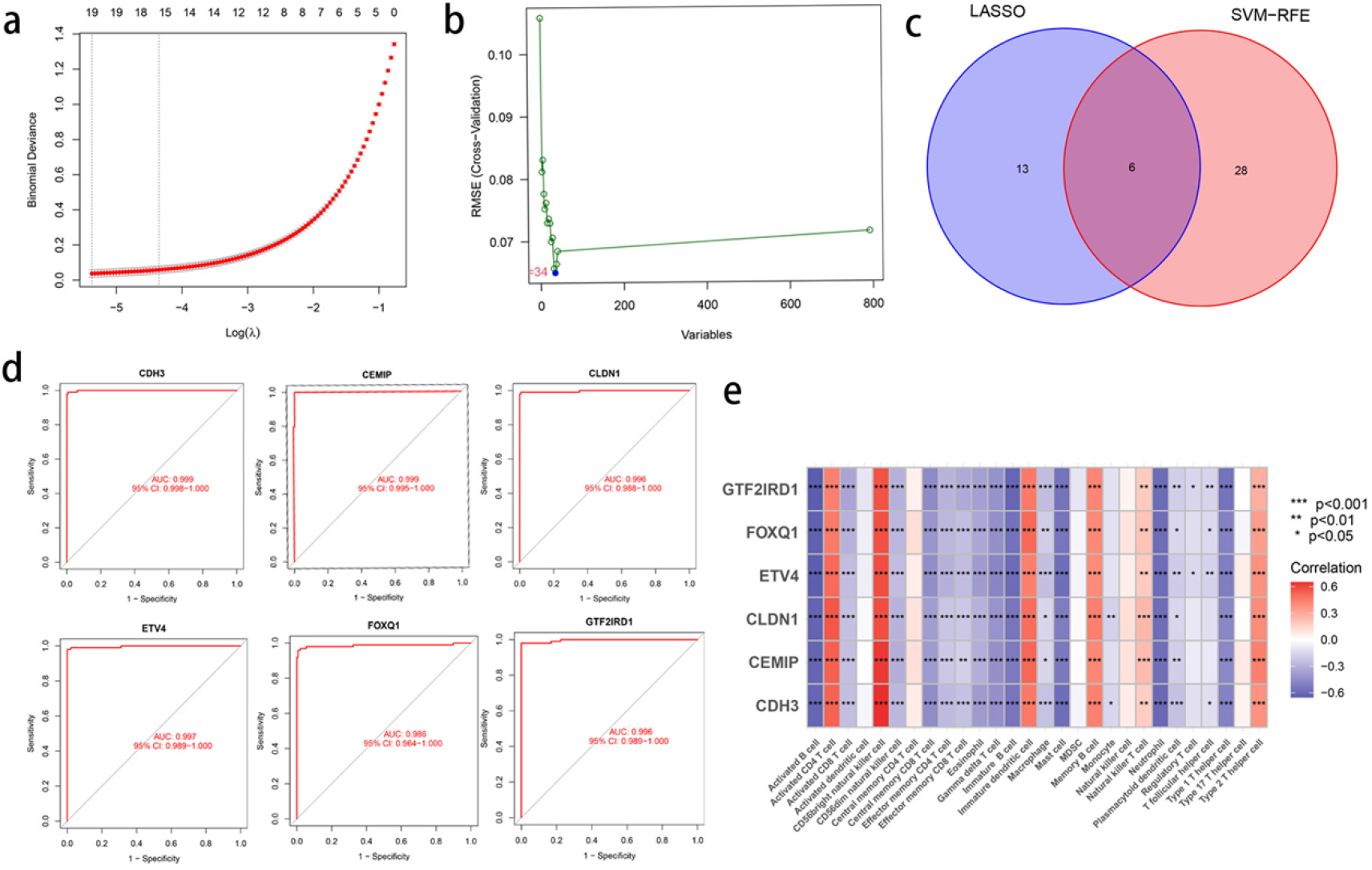
Finding feature genes by dimensionality reduction clustering via machine learning. (a) LASSO algorithm. (b) SVM-RFE algorithm. (c) Venn diagram showing the six key genes (CLDN1, CEMIP, ETV4, CDH3, GTF2IRD1, FOXQ1). (d) Diagnostic value of CLDN1, CEMIP, ETV4, CDH3, GTF2IRD1, and FOXQ1 in screening for MSS-type colorectal cancer. (e) ssGSEA analysis.

### 3.3: GTF2IRD1 is associated with poor prognosis in CRC patients

Our information had been acquired from the TCGA database and blanketed RNA-seq and clinical information from 521 colon cancer patients (27 solid normal tissue and 521 primary tumor tissue). Samples without clinical information or with less than 30 days of follow-up were excluded to minimize the interference of irrelevant factors. We covered the characterized genes screened above and carried out univariate COX regression evaluation to become aware of potential predictive genes(Fig 3a and b). Three genes have been recognized as prognostic genes. The three genes in the risk model had been recognized by lasso regression analysis. Prognostic markers for three genes have been developed using multivariate COX regression analysis, which includes GTF2IRD1, ETV4, and FOXQ1. Our risk rankings were calculated in the TCGA cohort based totally on their coefficients and the usage of the following formula. Risk rating = GTF2IRD1*expression stage (−0.036) + ETV4*(−0.010) expression stage + FOXQ1*0.013. Patients were categorized into high and low-risk groups based on the median patient risk score. Survival curves suggested that patients in the high-risk group had significantly lower OS than those in the low-risk group(Fig 3c). The ROC curves showed areas under the curve (AUC) of 0.555, 0.581, and 0.578 at 1, 3, and 5 years, respectively. (Fig 3d). We developed nomograms primarily based on different impartial prognostic elements to predict universal survival in sufferers with colorectal cancer(Fig 3g), which consequently validated the usage of calibration graphs(Fig 3h). Subsequently, we examined the patients’ clinicopathologic traits and threat rankings. Significant variations existed between the two corporations in the T4 and N0 stages and before graded tumors(Fig 3e and f).

**Fig 3.**
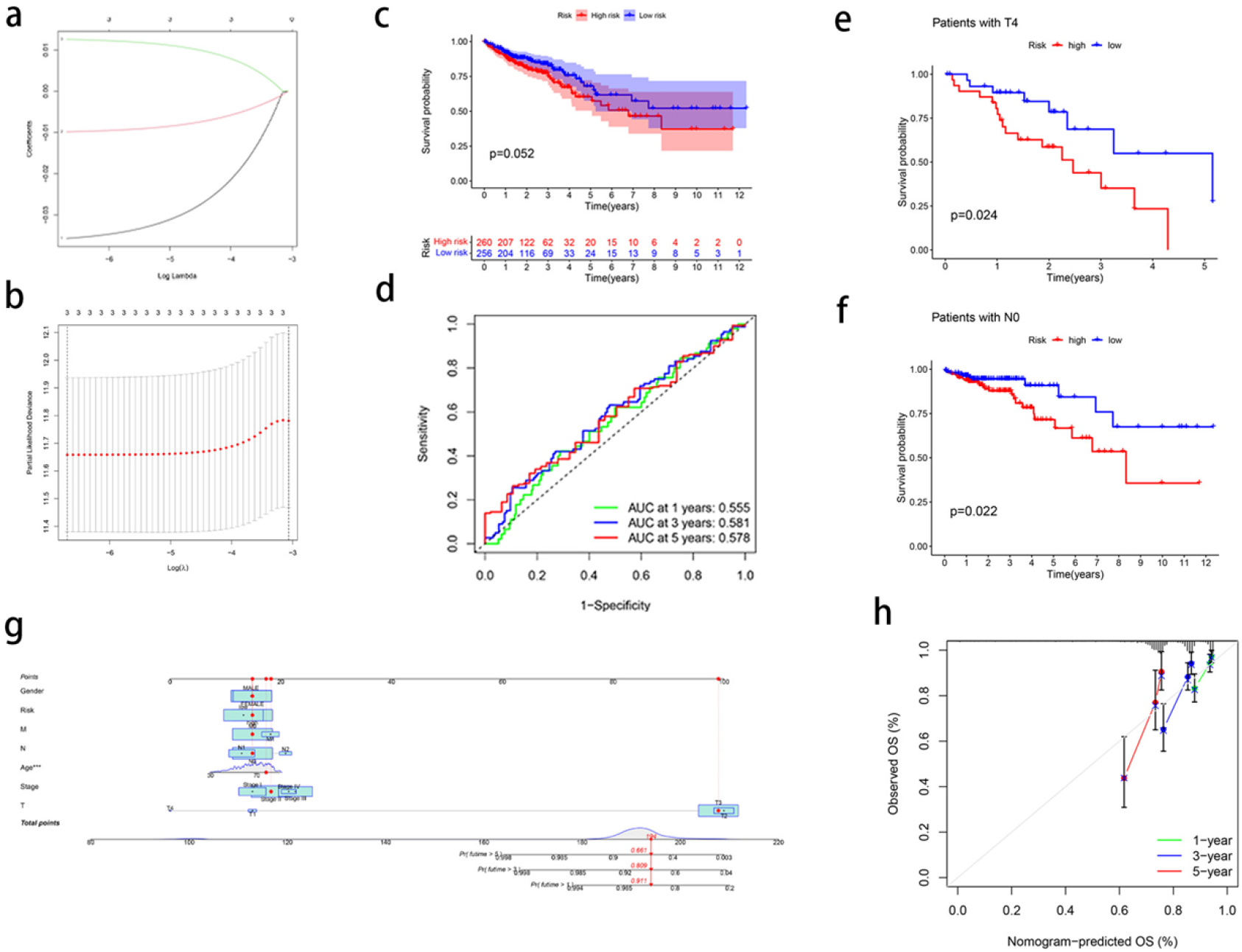
Establishment and validation of predictive models for colorectal cancer patients. (a) Cross-validation of three OS-related genes in LASSO regression. (b) Bias likelihood deviation of LASSO regression for three candidate genes. (c) Kaplan-Meier curves were used to analyze the OS of patients in two CRC risk groups. (d) ROC curves evaluated the predictive ability of the constructed risk models. (e-f) Relationship between risk scores and frequent clinicopathologic features. (g) Nomogram (h) Nomogram calibration curves.

### 3.4: GTF2IRD1 is highly expressed in MSS-type colorectal cancer and promotes its proliferation

We further analyzed these screened signature genes. First, we downloaded the GSE25071 dataset from the GEO database, removed the data of patients with MSS typing, and retained the data of patients with MSI typing and normal tissues. The results of principal component analysis showed significant principal component differences between the two groups(Fig 4a). Differential gene analysis revealed that GTF2IRD1 did not differ between the two groups and was a stable gene(Fig 4b). We also sought confirmation from the online database GEPIA2.0, and the results also matched the results we obtained(Fig 4c). Then, we confirmed the expression of GTF2IRD1 in two sorts of colorectal cancers at the mRNA and protein levels, respectively. QRT-qPCR and immunohistochemistry consequences confirmed that the expression of GTF2IRD1 used to be extended in MSS-type colorectal cancers, and there used to be No substantial distinction in the expression of MSI-type colorectal cancers(Fig 4d and g). HT29 cell line is usually used as a representative cell line for MSS-type colorectal cancers, and the LOVO cell line is used as a representative cell line for MSS-type colorectal cancers. We performed GTF2IRD1 knockdown on the HT29 cell line to verify the transfection efficiency and showed that SI-3 had the highest transfection efficiency(Fig 4e). We performed GTF2IRD1 knockdown on the HT29 cell line and LOVO cell line, respectively, with SI-3, followed by CCK-8 proliferation assay, which showed that the proliferation rate of the GTF2IRD1 knockdown group in the HT29 cell line was substantially decreased than that of the normal group. However, there was no considerable alternate in the proliferation rate of the GTF2IRD1 knockdown group in the LOVO cell line compared with that of the normal group(Fig 4f).

**Fig 4.**
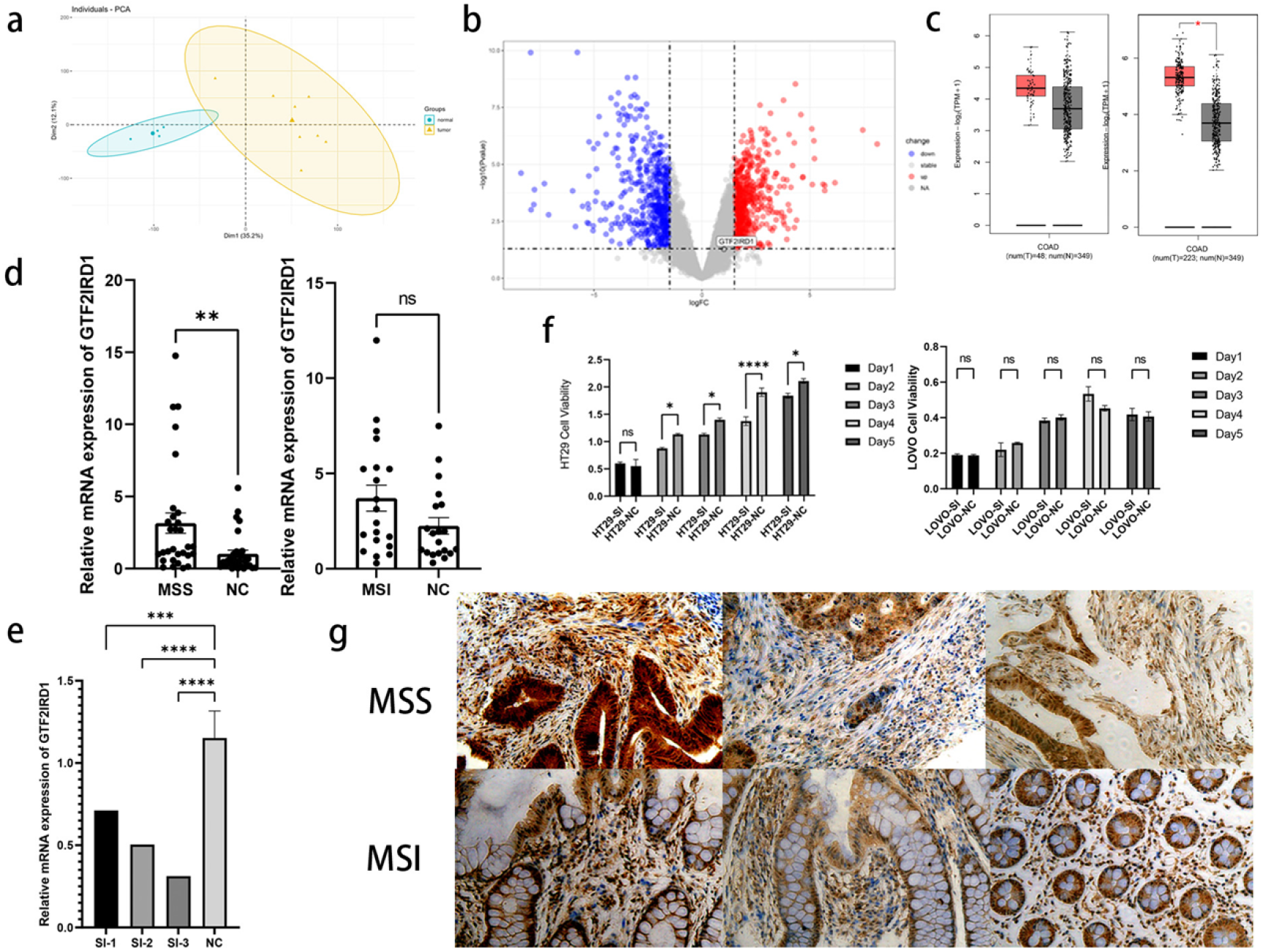
GTF2IRD1 is highly expressed and promotes proliferation in MSS-type colorectal cancer. (a) Principal component analysis of the two groups in GES 25071. (b) Differentially expressed genes between CRC specimens and normal specimens in the GSE 25071 dataset are shown in the volcano plot. (c) GEPIA 2.0 validation of GTF2IRD1 expression differences between MSI-type CRC and normal tissues on the left and between MSS-type CRC and normal tissues on the right. (d) Detection of GTF2IRD1 expression in MSI colorectal cancer tissues and MSS type colorectal cancer tissues by QRT-PCR. (e) QRT-qPCR of GTF2IRD1 mRNA expression in GTF2IRD1 siRNA-transfected HT29 cells, as well as control siRNA-transfected cells. (f) CCK-8 assay was used to measure the proliferation rate of HT29 cells and LOVO cells after transfection. (g) IHC detected differences in GTF2IRD1 protein expression between MSI-type CRCs and MSS-type CRCs. ns: not significant, *P < 0.05, **P < 0.01, ***P < 0.005, ****P < 0.0005.

## 4. Discussion

In this study, we first used bioinformatics to find the different genes between MSS-type colorectal cancer and normal tissues. We then used two machine learning algorithms to perform dimensionality reduction clustering on these genes, from which we found characteristic differential genes. Among these characterized differential genes, we further found GTF2IRD1, a gene highly expressed only in MSS-type colorectal cancer but stable in MSI typing, and explored their clinical relevance. We then experimentally verified the expression and function of this gene in the two bowel cancer types and demonstrated that this gene can promote the proliferation of MSS-type colorectal cancer and affect the prognosis of patients(14).

GTF2IRD1 is a member of the GTF21 gene family and encodes a team of multifunctional transcription factors(15). This gene has been studied in the field of oncology, including breast cancer(16), lung cancer, and pancreatic cancer. Previous studies have demonstrated the pro-carcinogenic effects of GTF2IRD1 in colorectal cancer, and GTF2IRD1 down-regulates the expression of the transforming growth factor-β receptor 2 (TGFβR2) in Smad4 mutant CRC(17). So, GTF2IRD1 is a conceivable oncogene and a biomarker with a terrible prognosis in human cancers, together with CRC.

The consequences of the KEGG enrichment analysis confirmed that the up-regulated differential genes had been generally enriched in the IL-17 signaling pathway, cell cycle, and different associated pathways, and the outcomes of the GSEA enrichment analysis confirmed that the up-regulated genes of MSS were generally enriched in the CELL cycle and CHEMOKINE_SIGNALING_PATHWAY and different related pathways. The outcomes of the GSEA enrichment evaluation confirmed that MSS up-regulated genes have been specially enriched in the CELL cycle and CHEMOKINE_SIGNALING_PATHWAY pathways. A previous article reported that the expression of GTF2IRD1 was positively correlated with the expression of genes associated to cell cycle progression and genes associated with the non-canonical Wnt-calcium pathway, which is known to regulate migration. Our enrichment analysis results matched the results reported in this article. ssGSEA results showed that GTF2IRD1 showed a significant negative correlation with most immune cells(18), such as Activated CD8 T cells, Effector memory CD8 T cells, etc. A previous article demonstrated that GTF2IRD1 promotes cell cycle progression by down-regulating TGFβR2 in CRC, and a recent article showed that up-regulation of transforming growth factor-β1 expression inhibited the activation of memory CD8+ T cells and promoted immune evasion(19). We speculate that GTF2IRD1 may also affect the cell cycle and immune cell infiltration by influencing the transforming growth factor-β signaling pathway. Still, the specific regulatory mechanism is not clear.

The application of machine learning in cancer research has the potential to transform our understanding of the disease and improve patient prognosis through early identification and more personalized treatment choices. In this study, we identified three prognostically relevant genes, GTF2IRD1, ETV4, and FOXQ1; KM survival curve results were not statistically significant, possibly because we included too small a sample size, resulting in insignificant differences between groups. We did not differentiate the included TCGA patients from microsatellite typing, so this point may affect the significance of the difference between the groups because our study proved that GTF2IRD1 was highly expressed in MSS typing and stable in MSI(20). So, although the results were not statistically significant, they may still be necessary in practical applications.

This study demonstrated that GTF2IRD1 expression was elevated in MSS-type colorectal cancers. We proved that this gene could promote the proliferation of MSS-type colorectal cancers and was related with bad prognosis in patients. The limitation of this study is that we hypothesized that GTF2IRD1 affects patients’ prognosis by regulating the cell cycle and down-regulating immune cell infiltration in the immune microenvironment of tumors. We still have yet to validate the specific mechanism further. We also did not explore the downstream pathways of GTF2IRD1.

## Acknowledgment

We want to thank the Department of General Surgery, General Hospital of Northern Theater Command (General Hospital of Shenyang Military Command) staff for providing us with access to the resources and facilities necessary to complete this research.

## Notes

### Competing Interest Statement

The authors have declared no competing interest.

## References

1. Wang Q, Zhang YF, Li CL, Wang Y, Wu L, Wang XR, et al. Integrating scRNA-seq and bulk RNA-seq to characterize infiltrating cells in the colorectal cancer tumor microenvironment and construct molecular risk models. Aging (Albany NY). 2023;15. doi: 10.18632/aging.205263. PubMed PMID: 38054820.

2. Li S, Zhang N, Yang Y, Liu T. Transcriptionally activates CCL28 expression to inhibit M2 polarization of macrophages and prevent immune escape in colorectal cancer cells. Transl Oncol. 2023;40:101842. doi: 10.1016/j.tranon.2023.101842. PubMed PMID: 38035446.

3. Cai L, Chen A, Tang D. A new strategy for immunotherapy of microsatellite-stable (MSS)-type advanced colorectal cancer: Multi-pathway combination therapy with PD-1/PD-L1 inhibitors. Immunology. 2024. doi: 10.1111/imm.13785. PubMed PMID: 38517066.

4. Goel A, Boland CR. Epigenetics of colorectal cancer. Gastroenterology. 2012;143(6):1442–60.e1. doi: 10.1053/j.gastro.2012.09.032. PubMed PMID: 23000599.

5. André T, Shiu KK, Kim TW, Jensen BV, Jensen LH, Punt C, et al. Pembrolizumab in Microsatellite-Instability-High Advanced Colorectal Cancer. N Engl J Med. 2020;383(23):2207–18. doi: 10.1056/NEJMoa2017699. PubMed PMID: 33264544.

6. Diaz LA L. DT. PD-1 Blockade in Tumors with Mismatch-Repair Deficiency. N Engl J Med. 2015;373(20):1979. doi: 10.1056/NEJMc1510353. PubMed PMID: 26559582.

7. Thibaudin M, Fumet JD, Chibaudel B, Bennouna J, Borg C, Martin-Babau J, et al. First-line durvalumab and tremelimumab with chemotherapy in RAS-mutated metastatic colorectal cancer: a phase 1b/2 trial. Nat Med. 2023;29(8):2087–98. doi: 10.1038/s41591-023-02497-z. PubMed PMID: 37563240.

8. Zhuang H, Zhang C, Hou B. GTF2IRD1 overexpression promotes tumor progression and correlates with less CD8+ T cells infiltration in pancreatic cancer. Biosci Rep. 2020;40(9). doi: 10.1042/bsr20202150. PubMed PMID: 32936232.

9. Yi S, Zhang C, Li M, Qu T, Wang J. Machine learning and experiments identifies SPINK1 as a candidate diagnostic and prognostic biomarker for hepatocellular carcinoma. Discov Oncol. 2023;14(1):231. doi: 10.1007/s12672-023-00849-2. PubMed PMID: 38093163.

10. Hamid MA, Pammer LM, Lentner TK, Doleschal B, Gruber R, Kocher F, et al. Immunotherapy for Microsatellite-Stable Metastatic Colorectal Cancer: Can we close the Gap between Potential and Practice? Curr Oncol Rep. 2024. doi: 10.1007/s11912-024-01583-w. PubMed PMID: 39080202.

11. Ding K, Mou P, Wang Z, Liu S, Liu J, Lu H, et al. The next bastion to be conquered in immunotherapy: microsatellite stable colorectal cancer. Front Immunol. 2023;14:1298524. doi: 10.3389/fimmu.2023.1298524. PubMed PMID: 38187388.

12. Fu X, Huang J, Zhu J, Fan X, Wang C, Deng W, et al. Prognosis and immunotherapy efficacy in dMMR&MSS colorectal cancer patients and an MSI status predicting model. Int J Cancer. 2024. doi: 10.1002/ijc.34946. PubMed PMID: 38594805.

13. Zhang L, Deng Y, Yang J, Deng W, Li L. Neurotransmitter receptor-related gene signature as potential prognostic and therapeutic biomarkers in colorectal cancer. Front Cell Dev Biol. 2023;11:1202193. doi: 10.3389/fcell.2023.1202193. PubMed PMID: 38099288.

14. Bras-Gonçalves RA, Rosty C, Laurent-Puig P, Soulié P, Dutrillaux B, Poupon MF. Sensitivity to CPT-11 of xenografted human colorectal cancers as a function of microsatellite instability and p53 status. British Journal of Cancer. 2000;82(4):913–23. doi: 10.1054/bjoc.1999.1019.

15. Nambara S, Masuda T, Kobayashi Y, Sato K, Tobo T, Koike K, et al. GTF2IRD1 on chromosome 7 is a novel oncogene regulating the tumor-suppressor gene TGFβR2 in colorectal cancer. Cancer Sci. 2020;111(2):343–55. doi: 10.1111/cas.14248. PubMed PMID: 31758608.

16. Huo Y, Su T, Cai Q, Macara IG. An In Vivo Gain-of-Function Screen Identifies the Williams-Beuren Syndrome Gene GTF2IRD1 as a Mammary Tumor Promoter. Cell Rep. 2016;15(10):2089–96. doi: 10.1016/j.celrep.2016.05.011. PubMed PMID: 27239038.

17. Plewa N, Poncette L, Blankenstein T. Generation of TGFβR2(−1) neoantigen-specific HLA-DR4-restricted T cell receptors for cancer therapy. Journal for ImmunoTherapy of Cancer. 2023;11(2). doi: 10.1136/jitc-2022-006001.

18. Fu J, Jin X, Chen W, Chen Z, Wu P, Xiao W, et al. Identification of the molecular characteristics associated with microsatellite status of colorectal cancer patients for the clinical application of immunotherapy. Frontiers in Pharmacology. 2023;14. doi: 10.3389/fphar.2023.1083449.

19. Borràs DM, Verbandt S, Ausserhofer M, Sturm G, Lim J, Verge GA, et al. Single cell dynamics of tumor specificity vs bystander activity in CD8+ T cells define the diverse immune landscapes in colorectal cancer. Cell Discovery. 2023;9(1). doi: 10.1038/s41421-023-00605-4.

20. Cao Q, Dan Z, Hou N, Yan L, Yuan X, Lu H, et al. Discovery and validation of colorectal cancer tissue-specific methylation markers: a dual-center retrospective cohort study. Clin Epigenetics. 2024;16(1):122. doi: 10.1186/s13148-024-01735-6. PubMed PMID: 39244604.

